# TerrestrialMetagenomeDB: a public repository of curated and standardized metadata for terrestrial metagenomes

**DOI:** 10.1101/796441

**Authors:** Felipe Borim Corrêa, João Pedro Saraiva, Peter F. Stadler, Ulisses Nunes da Rocha

**Affiliations:** Department of Environmental Microbiology, UFZ-Helmholtz Centre for Environmental Research, Leipzig, Saxony, 04318, Germany; Department of Computer Science and Interdisciplinary Center of Bioinformatics, University of Leipzig, Leipzig, Saxony, 04107, Germany

## Abstract

Microbiome studies focused on the genetic potential of microbial communities (metagenomics) became standard within microbial ecology. MG-RAST and the Sequence Read Archive (SRA), the two main metagenome repositories, contain over 202 858 public available metagenomes and this number has increased exponentially. However, mining databases can be challenging due to misannotated, misleading and decentralized data. The main goal of TerrestrialMetagenomeDB is to make it easier for scientists to find terrestrial metagenomes of interest that could be compared with novel datasets in meta-analyses. We defined terrestrial metagenomes as those that do not belong to marine environments. Further, we curated the database using text mining to assign potential descriptive keywords that better contextualize environmental aspects of terrestrial metagenomes, such as biomes and materials. TerrestrialMetagenomeDB release 1.0 includes 15 194 terrestrial metagenomes from SRA and MG-RAST. Together, the downloadable data amounts to 68 Tbp. In total, 199 terrestrial terms were divided into 14 categories. These metagenomes span 84 countries, 31 biomes and 7 main source materials. The TerrestrialMetagenomeDB is publicly available at https://webapp.ufz.de/tmdb.

## INTRODUCTION

A metagenome, in microbiome research, encompasses the genetic potential of a microbial community obtained through shotgun sequencing of DNA extracted from a sample (1). A few databases provide permanent storage and public access to DNA sequencing. The major database with these characteristics is the Sequencing Read Archive (SRA) (2). SRA is part of the International Nucleotide Sequence Database Collaboration (3) along with the European Nucleotide Archive (4) and the DNA Data Bank of Japan (5). Another important repository is MG-RAST (6), which also provides analysis services. Due to crescent availability of metagenomes in public databases, scientists are able to revisit publicly available data and to answer new hypothesis and research questions by applying recent or novel bioinformatic techniques. Reanalysis of public available data may lead to novel discoveries and insights, especially when data analyses of multiple studies are combined. For example, a study by Parks and collaborators (7) substantially expands the tree of life with the recovery of nearly 8 000 assembled genomes from 1 550 metagenomes and a meta-analysis study with 9 428 metagenomes defines the core species inhabiting the human gut (8). However, mining metagenomes of interest from databases is often not an easy task. For instance, to retrieve metagenomes from SRA can be very laborious since submitters often mislabel their datasets, making it impossible to distinguish metagenomes from amplicon sequencing data (9). The first initiative to suggest standards for metagenome submission was made in 2008 by the Genomic Standards Consortium (10). Soon after, these standards were implemented by MG-RAST (11). In 2011, SRA integrated BioProject and BioSample in their database (12), so that any submitted metagenomic sample must include the minimum information about a metagenome (13). To date, no studies have been published regarding the number of mislabelled or when no data was added, particularly regarding sample-related attributes. More recently, specialists started to improve and to curate standards within their fields. For example, Bernstein and collaborators (14) and Pasoli and collaborators (15) curated and standardized data from human-specific samples deposited in SRA. Nevertheless, prior TerrestrialMetagenomeDB, a resource focused on metagenomes obtained from terrestrial environments was not available. Here, we define terrestrial metagenomes as any environmental metagenome that belongs to terrestrial biomes as defined by Buttigieg and collaborators (i.e., ENVO:00000446) (16). In 2018, a resource called MGnify, former EBI-Metagenomics (17), was introduced (https://www.ebi.ac.uk/metagenomics/). Among its functionalities, MGnify can be used for the discovery of metagenomes from SRA. However, only for metagenomes that were analysed by this platform.

We created the TerrestrialMetagenomeDB, the first metadata database focused on terrestrial metagenomes, to help scientists researching terrestrial environments find metagenomes of interest that could be compared with novel datasets in meta-analysis studies. Our database consists of metadata related to biological samples and metadata describing technical aspects of the sequencing data. While the sample metadata supports biological questions in meta-analysis, the sequencing metadata can be crucial for bioinformatics. TerrestrialMetagenomeDB is not meant to replace recent efforts of BioSamples database to standardize and curate data (18), but to promote the exploratory possibilities of terrestrial metagenomes in a user-friendly interface, and to encourage comparison of public available data. Our resource combines the two current main databases (SRA and MG-RAST) and provides manually curated metadata that could be useful in meta-analysis studies.

## MATERIAL AND METHODS

### Database construction

The TerrestrialMetagenomeDB was constructed as follows. Briefly, we retrieved metadata of metagenomes from the source databases and parsed and standardized sample attributes. Next, we identified terrestrial metagenomes and removed marine samples. Finally, we combined the metadata of terrestrial metagenomes of SRA and MG-RAST, removed potentially entries that were non-metagenomes (targeted approaches and genome sequencing) and did not belong to terrestrial biomes and implemented the web application. A summarized graphic representation of the construction and availability of the TerrestrialMetagenomeDB is depicted in Figure 1.

**Figure 1.**
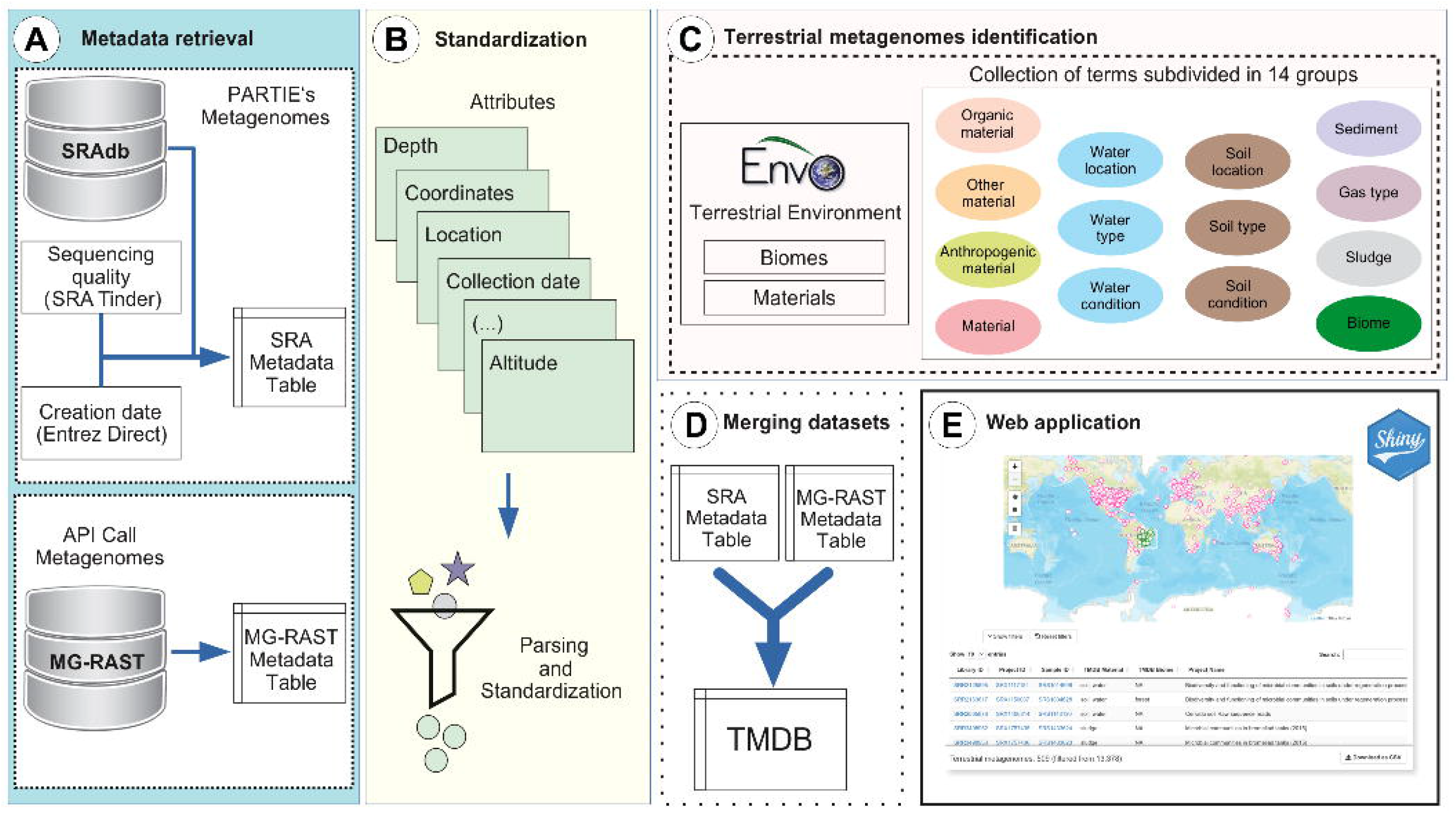
Overview of the TerrestrialMetagenomeDB (TMDB) construction and availability. The construction of TMDB comprises: (A) Metadata retrieval for metagenomes present in SRA and MG-RAST; (B) Standardization of attributes; (C) Identification of terrestrial metagenomes; and (D) merging of SRA and MG-RAST metadata. (E) The TMDB was made available through a user friendly Shiny web application.

#### Data retrieval

We retrieved the data for TerrestrialMetagenomeDB from SRA and MG-RAST as they are the largest repositories of publicly available metagenomes. For SRA, we selected identifiers of libraries (SRA Runs) assigned as metagenomes by PARTIE (9), a tool that periodically checks if libraries are correctly annotated as metagenomes. The complete list of identifiers is available at PARTIE’s Github (https://github.com/linsalrob/partie) as ‘SRA_Metagenome_Types.tsv’. After, we retrieved the respective metadata using the SRAdb R package (19), which provides local access to all metadata entries from SRA. Additionally, we retrieved quality scores of the sequences with the tool SRA-Tinder (https://github.com/NCBI-Hackathons/SRA_Tinder). We also retrieved the date of creation of the libraries using Entrez Direct (https://www.ncbi.nlm.nih.gov/books/NBK179288), because these dates are not available through SRAdb. To avoid potential non-WGS datasets we filtered out entries where ‘library_selection’ was filled with ‘PCR’ or ‘library_strategy’ was filled “AMPLICON”. For MG-RAST, all entries were requested to its application program interface (API). To select only metagenomes at MG-RAST, we filtered the entries’ metadata where values of ‘investigation_type’ and ‘seq_meth’ were equal to ‘metagenome’ and ‘WGS’ for whole genome sequencing.

#### Standardization of attributes

In SRAdb, all the sample attributes are available in a single field and the attribute names are written in many different ways. For these reasons, we standardized synonyms and screened attribute names and their respective values. For SRA, we standardized nine different attributes: sample latitude, sample longitude, sample depth, sample elevation, sample altitude, sample temperature, sample pH, sample location and sample collection date. Those attributes were parsed from the SRAdb field named ‘sample_attribute’. We removed parsed attributes with less than 10 occurrences. Further, we grouped the remaining attributes by synonyms (Supplementary Table S1). Coordinates were standardized to the format of Decimal Degrees (round to 6 decimal digits, resolving up to 0.11 m). Dates were standardized to the international standard according to ISO 8601 (YYYY-MM-DD) and dates that were not between 1950 and the current year were marked as “NA”. Countries where the samples were collected were manually labelled according to country names of standard ISO 3166−1. For MG-RAST, the equivalent attributes were already available (except sample pH) from the API retrieval and, when necessary, they were adapted to the above mentioned formats. Additionally, we distinguished between datasets containing assemblies and those containing sequencing reads by adding an attribute named “average_length” (basepairs count / sequences count) per metagenome. From the average length, we inferred assembled Illumina and 454 datasets and added annotation in the attribute named “assembled”. We annotated as “Yes” (i.e., assembled data) when the average length was greater than 600 bp.

#### Identification of terrestrial metagenomes

A set of words of ‘Environmental Material’ and ‘Terrestrial Environment Biome’ was adapted from The Environment Ontology (ENVO) (16). To select the most relevant words, every word was queried against the collected metadata, and the relevant words were grouped. After, we added the prefix ‘TMDB’ to tag our terrestrial groups (Supplementary Table S2). Further, we queried the words in the complete metadata and assigned them to each terrestrial group. Metagenomes were classified as terrestrial when at least one terrestrial word was present in the metadata.

#### Removal of marine samples

We used two different approaches to remove marine samples. These different approaches were based on the presence or absence of coordinates for each metagenome. For metagenomes with coordinates, we removed entries with coordinates outside land boundaries (i.e., in the sea). To that end we used ‘is-sea’ (https://github.com/simonepri/is-sea). For metagenomes without coordinates available, we searched in the metadata for terms that indicated the given metagenomes are potentially from the marine environment. The terms were respectively ‘sea’, ‘marine’ and ‘ocean’.

#### Combining SRA and MG-RAST metadata

We selected a collection of equivalent and comparable attributes present in both databases and combined those into the metadata found in the TerrestrialMetagenomeDB (Supplementary Table S3). Only three attributes related to library sequencing quality scores were unique and specific to SRA or MG-RAST; respectively, ‘quality_above_30_SRA’, ‘mean_quality_SRA’, and ‘drisee_score_raw_MGRAST’.

#### Removal of non-terrestrial and non-metagenomic datasets

From the final combined metadata, we filtered out datasets related to human-derived samples by searching for terms like “human” and “homo-sapiens”. Likewise, for filtering out non-metagenomic datasets we used keywords related to amplicon sequencing, genome sequencing and genome assembly. A list with the regular expressions used to perform the filtering is listed in the Supplementary Table S4.

#### Web app implementation

TerrestrialMetagenomeDB web-interface was implemented using Shiny (version 1.3.2) for R (version 3.4.2). The map in the ‘Interactive map’ tab was added using the *leaflet* package (version 2.0.2), and the selection toolbox was created with the *leaflet.extras* package (version 1.0.0). The function for selecting points on the map was built using the *geoshaper* package (version 0.1.0) and the *sp* package (version 1.3-1). The ‘Interactive map’ and ‘Complete dataset’ data tables were set using the *DT* package (version 0.7). The ‘Help’ and ‘Contact’ R markdown texts were created with the *markdown* package (version 1.0) and the *knitr* package (version 1.23). The modal that opens with the ‘Interactive map’ was generated with the package *shinyBS* (version 0.61).The mouse over tooltips were generated with the package *shinyBS* and the numeric range inputs were implemented with the package *shinyWidgets* (version 0.4.8).

## RESULTS

### Database content

The current manuscript describes the TerrestrialMetagenomeDB release 1.0, where 15 194 metagenomes from the terrestrial environment are available. Those cover 11 years of experiments, since the first terrestrial metagenome was submitted to the SRA on May 2008 and the latest registered at MR-RAST on May 2019. Among those, 6 845 metagenomes are derived from SRA (45%) and 8 349 from MG-RAST (55%). Since SRA and MG-RAST are independent from each other, identical datasets can exist in both databases. In addition, metadata provided by submitters was insufficient to determine which datasets overlap. According to the location where the samples were collected, metagenomes span all 7 continents and 84 countries. Most of the libraries available were sequenced with Illumina sequencing technologies (86%), followed by LS454 (6%), Ion Torrent (2%) and others (6%) (Figure 2).

**Figure 2.**
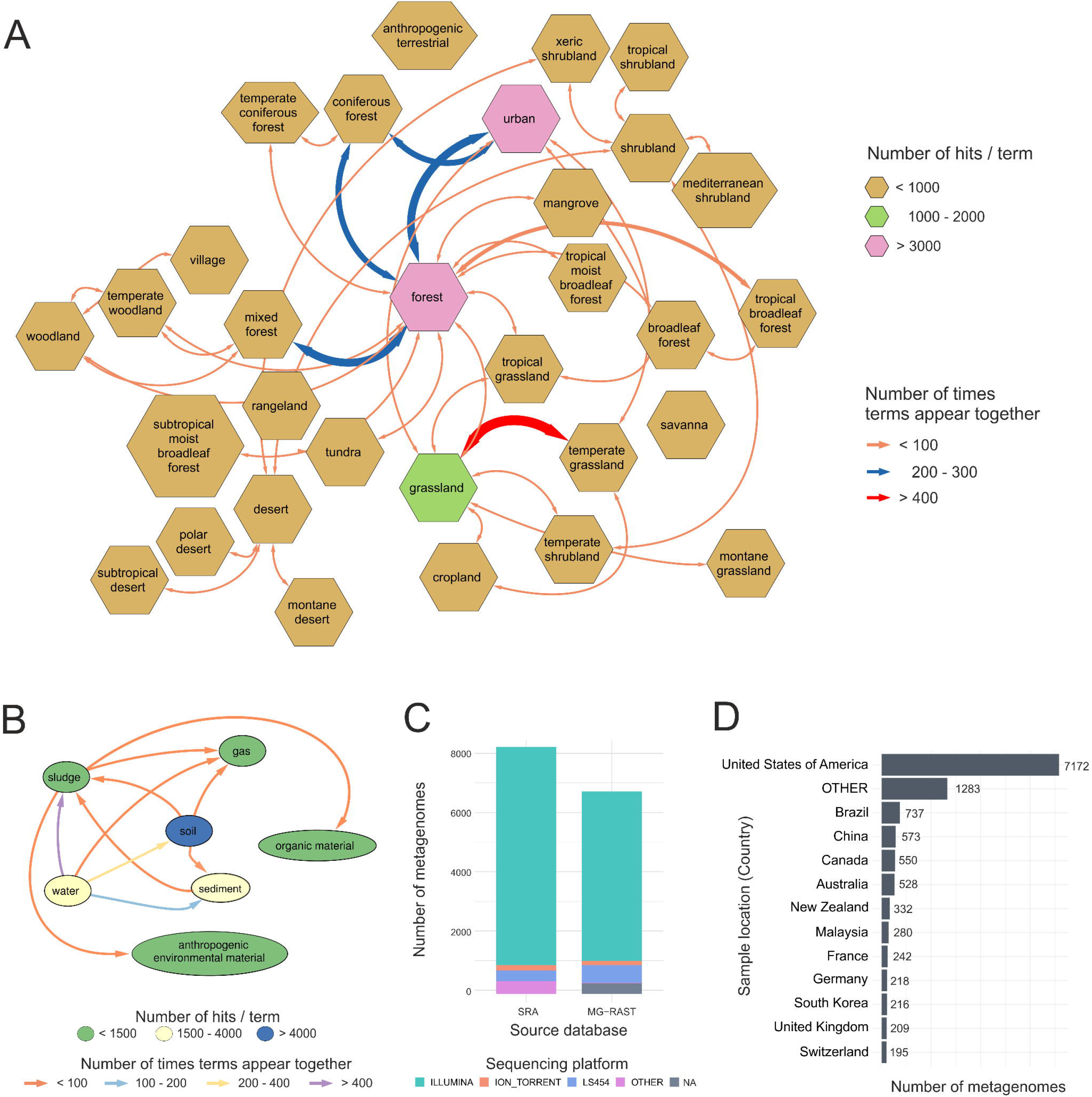
Descriptive statistics of the TerrestrialMetagenomeDB content. (A) Network representation of the frequencies of ‘biome’-related terms in the database (polygon shape). The frequencies of pairs of ‘biome’ terms found in the database are represented by coloured arrows. (B) Network representation of the frequencies of ‘material’-related terms in the database (ellipse shape). The frequencies of pairs of ‘material’ terms found in the database are represented by coloured arrows. (C) Bar plot showing the distribution of the country of origin of the metagenomic samples (Sample location), in this plot the not assigned values (NA‘s) were omitted. (D) Bar plot of the distribution of sequencing technologies (Sequencing platform) per database of origin (Source database).

Regarding the quality of the library reads, 99% of the SRA metagenomes have quality scores. For MG-RAST metagenomes, another measurement of quality named DRISEE (13) is available, where 67% of the metagenomes have this attribute annotated.

#### Terrestrial metagenomes

The most populated ‘TMDB terrestrial attributes’ were ‘TMDB material’ with 76% of present values followed by ‘TMDB biome’ with 41% of values annotated with our pipeline. The top terms in ‘TMDB material’ were soil (6209), water (2655), sediment (1561), sludge (1244) and organic material (574). The top terms in ‘TMDB biome’ were urban (1881), forest (1623), grassland (1039), temperate grassland (475) and shrubland (319). Many terrestrial terms identified in the metadata co-occurred for the same metagenome, for example: the ‘TMDB Biome’ terms identified in the metagenomic library ‘mgm4819186.3’ were ‘forest’ and ‘urban’. The frequency and co-occurrence of terrestrial terms of ‘TMDB biome’ and ‘TMDB material’ present in the database can be visualized in Figure 2 A-B. The other 12 terrestrial categories defined in this work appeared in a lower frequency. From those, the top populated attributes were ‘TMDB organic material’ 12%) and ‘TMDB soil location’ (10%), and all the others were available in less than 5% of the metagenomes metadata. Supplementary Figure S1 depicts the percentage of missing values per attribute in the current TMDB data.

### Usage and functionalities

The TerrestrialMetagenomeDB user interface is divided in two main sections so users can choose the section that better fits their needs. In summary, the first section “Complete dataset” holds the full content of the databases’ current version. On the other hand, the “Interactive map” section provides a more intuitive way of selecting metagenomes directly from the world map, although being limited by the metagenomes with a pair of valid geographic coordinates available. To provide practical examples, we made 3 video tutorials about the usage of the web application. A link to the tutorials can be found in the item 1 of the “Help” tab in the web application.

#### Complete dataset

The ‘Complete dataset’ tab contains the sum of all entries with and without coordinates (15 194 in total). In this tab the initial data table is displayed with all the entries, what allows filtering and searching in the complete database. For filtering, a set of 6 filters is placed on top of the datatable for the most important attributes. By pushing the button “More filters”, the filters dashboard is expanded downwards to show all 33 filters. A panel is fixed on the bottom displaying the current number of filtered metagenomes, so users can keep track of how each filtering step is shaping the data. If no filter is applied, the whole dataset (the full TerrestrialMetagenomeDB data) can be downloaded as a CSV-file.

#### Interactive map

The ‘Interactive map’ tab allows users to interactively explore the world map and select metagenomes from all around the globe (Figure 3). Two drawing tools (rectangle and polygon) are available and allow easy selection of plotted points in the map by marking everything that is inside the selected boundaries. Individual points in the map may indicate several samples collected in the same coordinates. Therefore, we opted to only allow the selection of samples by using the drawing tool. This selection step can be performed multiple times in various regions of the map or re-started by clearing the drawn selection layers. Once the selection step is finished, all the marked points are displayed in the interactive data table below the map. This tab has exactly the same functionalities as described above, but without the filter for geographical coordinates, since this can be done directly on the map. Also, all 31 filters are hidden and can be shown by pushing the “Show filters” button. The columns ‘library id’, ‘project id’ and ‘sample id’ (when valid) are hyperlinked to the original source databases. A search box on the top-right of the data table allows the search for any text inside the selected metagenomes metadata. Reactively, the selected points in the map will be redrawn according to any filtering. The selected metadata (filtered or not) can be downloaded as a CSV-file.

**Figure 3.**
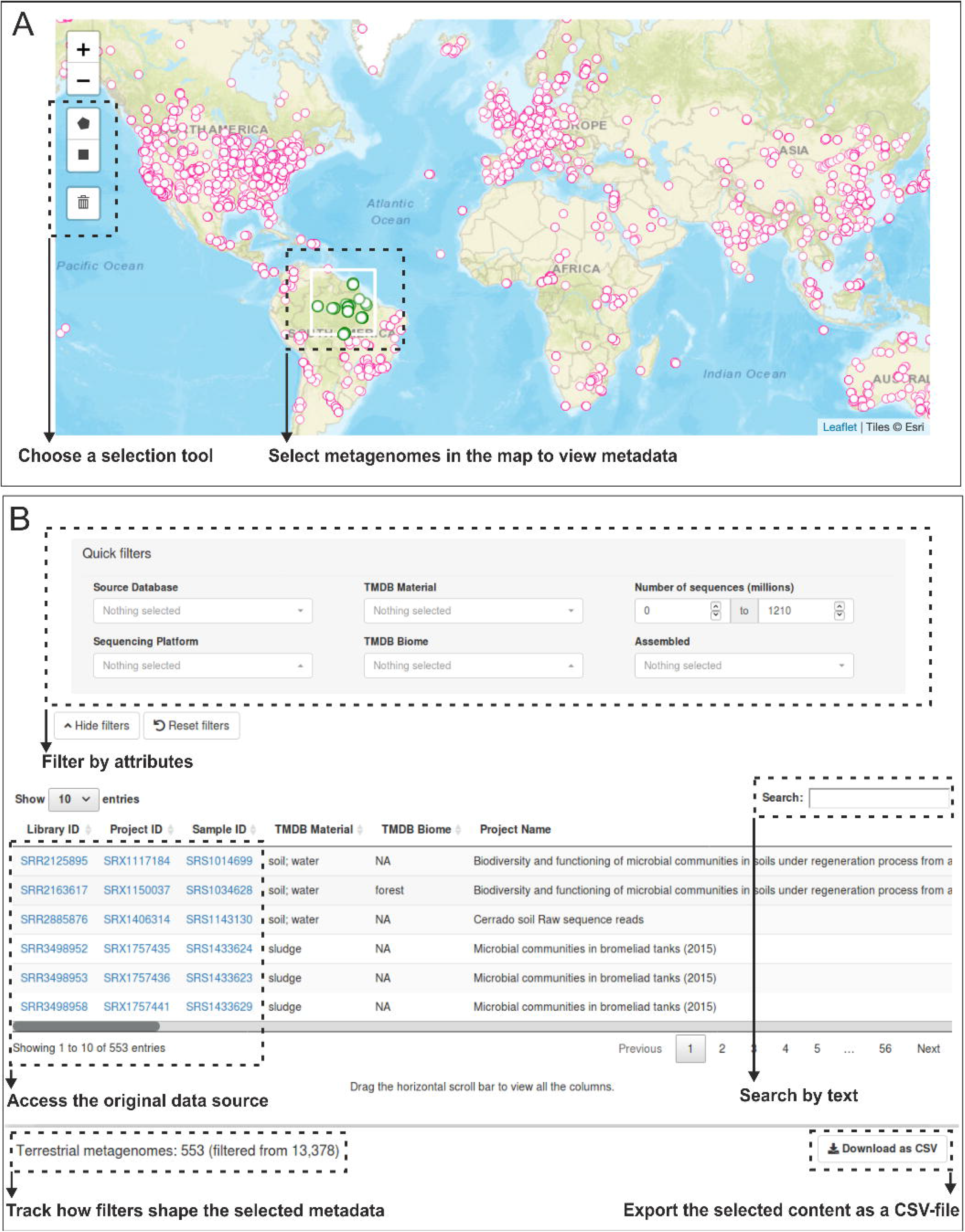
Overview of the TerrestrialMetagenomeDB user-interface. (A) Metagenomes can be selected in the ‘Interactive Map’ using a selection tool. (B) Metadata related to the selected entries is shown in the data table and can be further filtered and exported. For illustrative purposes, only the set of “Quick filters” is depicted.

#### Downloading metagenomes of interest

TerrestrialMetagenomeDB is not a repository of DNA sequence data. Once metagenomes are selected, they have to be downloaded from their respective repositories. A short guide on how to download the actual sequencing data from the original repositories can be found in the database user interfaces’ tab named ‘Help’. To facilitate the download, we provided a script called “tmdb_downloader.py” that takes as input the downloaded CSV-file from our database. Also, a video tutorial on how to use the script is available in the tutorials playlist.

#### Suggestion for good practices

To help scientists analyse their first metagenomes, a guide for ‘good practices’ when preparing the metadata for metagenomic studies can be found in the database user interfaces’ tab named ‘Help’, at the item 6 ‘What should I do to include my metagenomes in TMDB?’.

## CONCLUSION

TerrestrialMetagenomeDB is the first database to centralize and standardize metadata present at the Sequence Read Archive and MG-RAST for terrestrial metagenomes. We arranged terrestrial terms derived from the environment ontology ENVO with the help of scientists from different fields of terrestrial research and identified those terms both in MG-RAST and SRA.

TerrestrialMetagenomeDB is in its release 1.0 and it will get two updates per year, due to the exponential number of novel metagenomes added to public repositories. We believe that our database improves the current necessity of adequately described metadata (or contextual data) that will make possible querying and interpretation across projects and meta-analyses.

## AVAILABILITY

The TerrestrialMetagenomeDB is available at https://webapp.ufz.de/tmdb.

## ACKNOWLEDGEMENT

We thank Natascha Menezes Bergo for testing the TerrestrialMetagenomeDB web application and Rodolfo Brizola Toscan for creating the script to download the sequencing data. We are grateful to Dr. Sebastian Canzler, Dr. Andreas Schüttler, Dr. Matthias Bernt and Sven Petruschke for the support with the Shiny app deployment.

## FUNDING

This work was supported by the Helmholtz Association (Germany) through the Young Investigator Group [VH-NG-1248]. Funding for open access charge: Helmholtz Association.

## CONFLICT OF INTEREST

None declared.

## Notes

https://webapp.ufz.de/tmdb/

