## Supplementary figures and images for "TerrestrialMetagenomeDB: a public repository of curated and standardized metadata for terrestrial metagenomes"

### Suppl_Figure_S1.png

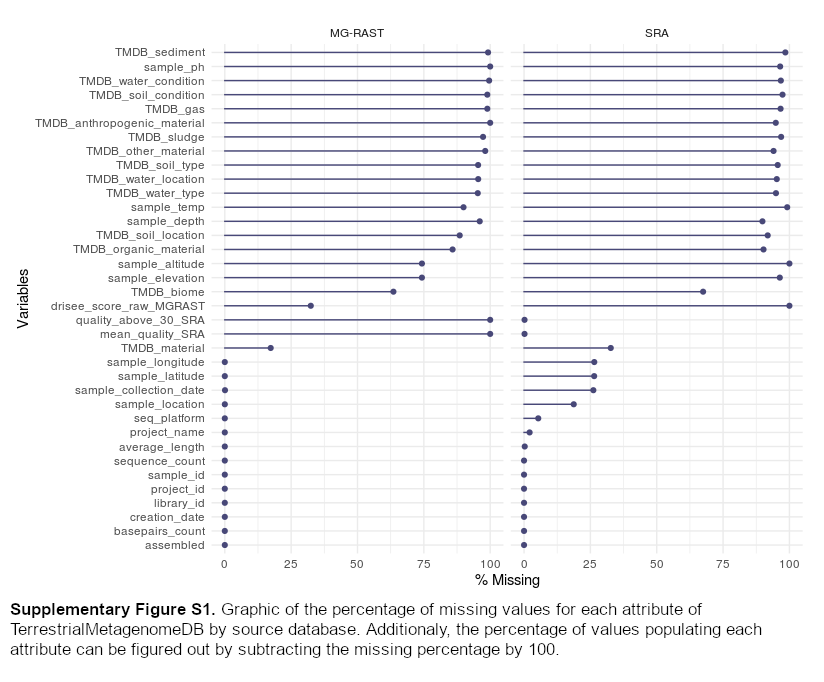
